# Transcriptome atlas of *Phalaenopsis equestris*

**DOI:** 10.1101/2020.11.03.366260

**Authors:** Anna V. Klepikova, Artem S. Kasianov, Margarita A. Ezhova, Aleksey A. Penin, Maria D. Logacheva

**Author notes:** Corresponding Author: Anna V. Klepikova ^1^, Bolshoy Karetny per. 19, build. 1, Moscow 127051, Russia.

## Abstract

The vast diversity of Orchidaceae together with sophisticated adaptations to pollinators and other unique features make this family an attractive model for evolutionary and functional studies. The sequenced genome of *Phalaenopsis equestris* facilitates Orchidaceae research. Here we present an RNA-seq based transcriptome map of *P. equestris* which covers 19 organs of the plant including leaves, roots, floral organs and shoot apical meristem. We demonstrated the high quality of the data and showed the similarity of *P. equestris* transcriptome map with gene expression atlases of other plants. The transcriptome map can be easily accessed through our database Transcriptome Variation Analysis (TraVA) visualizing gene expression profiles. As an example of the application we analyzed the expression of *Phalaenopsis* “orphan” genes – the ones that do not have recognizable similarity with genes of other plants. We found that about a half of them are not expressed; the ones that are expressed have a predominant expression pattern in reproductive structures.

## Introduction

The enormous diversity of orchids traditionally attracts attention of plant biologists. Orchidaceae comprises about 25 thousand of species, which makes it the largest plant taxon (Cai et al., 2015). The diversification of orchids has evolved along with complex pollinator-adapted flower structure (Cozzolino & Widmer, 2005), CAM-photosynthesis and epiphytism (Silvera et al., 2009).

Genome assembly of *Phalaenopsis equestris* (the horse phalaenopsis) (Cai et al., 2015) provided novel opportunities for evolutionary and functional studies of Orchidaceae. Genome assembly was used for the functional studies of transcription factor families (Lin et al., 2016; Valoroso et al., 2019), somatic embryogenesis (Chen et al., 2019), retrotransposon insertions (Hsu et al., 2019), as well as for evolutionary studies of ancient polyploidy (Barrett et al., 2019). However, transcriptome resources of *P. equestris* remain limited even though the de novo transcriptome assembly was performed based on RNA sequencing of 11 organs (Niu et al., 2016).

In our study we present a transcriptome map of *P. equestris* consisting of 19 samples in two biological replicates. High-quality RNA of orchid organs and tissues was sequenced using Illumina technology resulting in 1 687 M reads. We compared expression characteristics of *P. equestris* transcriptome map with gene expression atlases of other plants to provide evidence of reliability of our data. Transcriptome map of *P. equestris* can be applied in a great variety of functional studies.

## Materials & Methods

### Growing conditions

Plants were grown in a climate chamber under a 16 h light/8 h dark cycle at 22°C and 50–60% relative humidity. Samples were collected in two biological replicates; each replicate consists of at least seven plants. Sample collection was performed within two hours (Zeitgeber time ZT8-10) to reduce the influence of the circadian cycle.

### RNA extraction, library preparation and sequencing

RNA was extracted using the RNeasy mini kit (Qiagen, The Netherlands) following the manufacturer’s protocol. To ensure a high quality of *Phalaenopsis* samples, RNA was analyzed using capillary electrophoresis on Agilent Bioanalyzer 2100. cDNA libraries for Illumina sequencing were constructed using the NEBNext Ultra II RNA Library Prep Kit for Illumina (New England BioLabs, MA, USA) following the manufacturer’s protocol in 0.5 of the recommended volume (due to low RNA quantity in such samples as shoot apical meristem). cDNA libraries were sequenced with the HiSeq4000 and NextSeq500 (Illumina, CA, USA) instruments (50 bp and 75 bp single read run).

### Read mapping

Read trimming was performed using Trimmomatic version 0.36 (Bolger, Lohse & Usadel, 2014) in a single read mode and parameters “ILLUMINACLIP:common.adapters.file:2:30:10 LEADING:20 TRAILING:20 SLIDINGWINDOW:4:15 MINLEN:30”. For read mapping genome assembly and annotation of *P. equestris* from PLAZA database (version 4.5) was used. Trimmed reads mapped on the genome assembly using Spliced Transcripts Alignment to a Reference (STAR) version 2.4.2 (Dobin et al., 2013) in the “GeneCounts” mode and parameters “--sjdbOverhang 59 --sjdbGTFfeatureExon exon --sjdbGTFtagExonParentTranscript gene_id” to obtain counts of uniquely mapped reads on each gene.

### Expression characteristics of transcriptome map

Gene read counts obtained with STAR were normalized on library size using size factors, as described in (Anders & Huber, 2010). A threshold of five or higher normalized read counts in each biological replicate was used to define expressed genes.

To describe gene expression pattern Shannon entropy values were calculated for expressed in at least one sample genes (Schug et al., 2005). In order to avoid overrepresentation of certain plant organs, the samples were grouped using distances on clustering tree: gene expression levels were averaged if samples had distance (1 - Pearson r^2^) less than 0.1. Sample groups are listed in Table S1.

### Data availability

The RNA-seq raw data of transcriptome map were deposited in NCBI Sequence Read Archive (SRA) under BioProject accession PRJNA667255. The TraVA database can be accessed at http://travadb.org/browse/Species=Phalaenopsis_equestris/.

## Results

### Transcriptome map construction

Ornamental orchid *P. equestris* comprises three varieties and numerous hybrids of various flower colors and sizes (Hsu & Chen, 2016). To create transcriptome atlas we chose *P. equestris* var. blue (orchidee.su) as clonal plants are available for the cultivar which helps to reduce interindividual variability. We have collected 31 samples covering main plant organs and developmental stages such as roots, young and mature leaves, floral organs, flower buds, and meristems. Each sample was collected in two biological replicates, and each replicate was pooled from at least seven plants. Sample RNA was sequenced on Illumina platform resulting in 29 M - 65 M raw single reads (38 M median) for each sample (for sequencing statistics see Table S2). After removing low-quality reads and technical sequenced 98.7-99.8% of reads remained (Table S2).

Reads were mapped on the reference genome of *P. equestris* (Cai et al., 2015) with only one match allowed (unique mapping); 9.2-89.6% of high-quality reads were successfully mapped (Table S2). 12 samples showed extremely low percentage of read mapping; unmapped reads were identified as sequences belonging to Cymbidium mosaic virus (GenBank accession MK816927) which are known to persist in the majority of *P. equestris* population and affect mainly mature and senescent tissues (Koh, Lu & Chan, 2014). As the library size of infected samples was insufficient and can distort the conclusions we excluded samples with a percentage of mapped read lower than 35% in at least one biological replicate. The remained samples had 37.3-89.6% of uniquely mapped reads with median of 81.6%.

Thus, we constructed a transcriptome map of *P. equestris* covering 19 organs and parts of the plant. Floral organs (anthers, labellum, inner and outer tepals), leaves at different developmental stages, axes (inflorescence and pedicel), shoot apical and inflorescence meristems, and root parts were taken into analysis (for detailed description of samples see Table S3). The biological replicates showed high consistency (median Pearson r^2^ = 0.99, Table S4).

Clustering of samples generally reflects plant body plan and groups organs with similar morphology and physiology (Klepikova & Penin, 2019). Hierarchical clustering of *P. equestris* samples showed the same pattern (Fig. 1A). Sample clusters were formed by floral organs, leaf parts, meristems and young leaves, inflorescence internode and root; young and mature anthers were an outgroup for the other samples, similar to *A. thaliana,* rice, and maize (Nobuta et al., 2007; Wang et al., 2010; Stelpflug et al., 2016; Klepikova et al., 2016). The distances between samples on clustering tree were closer than in other species we observed (Klepikova et al., 2016; Penin et al., 2019), which can be explained by the lack of older tissues in *P. equestris* transcriptome atlas.

**Figure 1.**
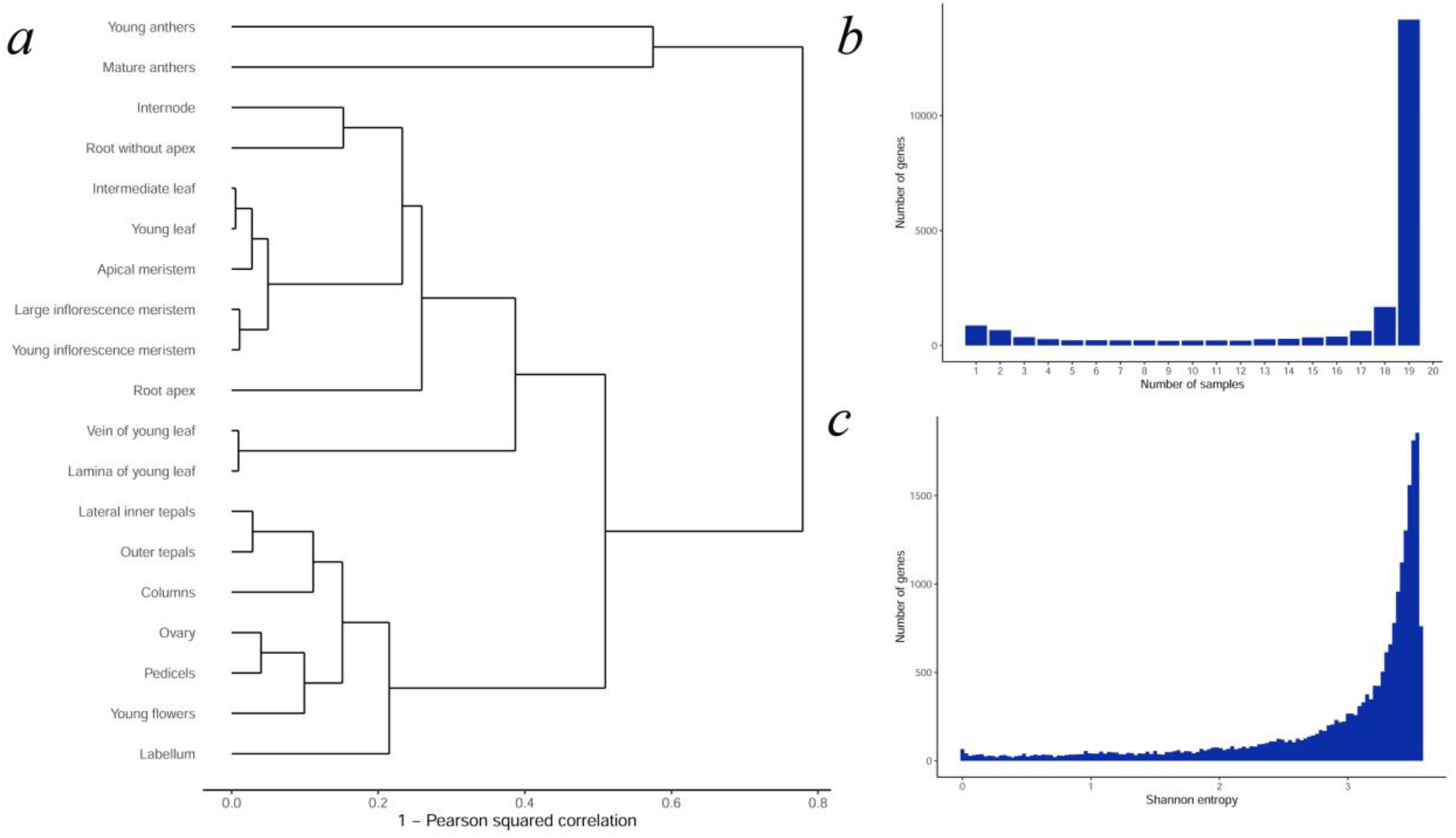
Expression characteristics of the P. equestris transcriptome map: (a) Hierarchical clustering tree of transcriptome map samples; (b) The distribution of genes by the number of samples where gene is expressed. Only expressed genes with 5 or more normalized read counts in each biological replicate were considered; (c) The distribution of Shannon entropy of P. equestris genes.

We compare our samples with publicly available *P. equestris* transcriptomes (Table S5). In general, the clustering of samples was consistent (Fig. S1), though leaf and column from the BioProject PRJNA288388 (Niu et al., 2016) form outgroup to all other samples.

### Expression characteristics of *P. equestris*

*Phalaenopsis* genome annotation (PLAZA database, version 4.5) includes 29 431 protein-coding genes. Among them 14 174 (48%) genes were expressed in all samples (using five reads in each biological replicate as a threshold), when transcripts of 21 671 (74%) genes were found in at least one sample. These values are in the range of typical expressed gene numbers across plant transcriptome maps (Klepikova & Penin, 2019). As in other species, samples demonstrated similarity in the number of expressed genes, which varied form 15 612 (53%) in shoot apical meristem to 18 947 (64%) in ovules before pollination (Table S6).

### Expression patterns of *P. equestris* genes

The study of gene expression pattern can shed light on the biological function of the gene and place it among essential for a plant existence ubiquitously expressed genes or precise regulators of tissue features – sample-specific expressed genes. We used two approaches to define gene expression patterns. A number of samples where gene is expressed is the simplest method to characterize expression pattern width, as was shown for *Nicotiana tabacum* (Edwards et al., 2010) or *Vigna unguiculata* (Yao et al., 2016). The majority of genes (16 486) were expressed in 17 or more samples; the second peak (1 896 genes) of the distribution is formed by genes expressed in 3 or less samples (Fig. 1B). The main patterns of tissue-specific genes were anthers (56% of tissue-specific genes), roots (11%), and meristems (both shoot apical and inflorescence meristem, 8%). The high number of anther-associated genes are known for *A. thaliana* (Klepikova et al., 2016) and is expected for *P. equestris* as young and mature anthers are the most distant samples on clustering tree (Fig. 1A).

While useful, such approach depends on an arbitrary threshold which separates expressed and non-expressed genes and does not take into account the variation of expression level between samples. To overcome the issue, we used Shannon entropy as a measure of expression pattern width: low entropy values correspond to tissue-specific genes, while high values mark ubiquitously expressed genes (Schug et al., 2005). The distribution of Shannon entropy in *P. equestris* was significantly skewed to the right revealing major part of wide-expressed genes (Fig. 1C) similarly to *A. thaliana, Solanum licopersicum* and *Zea mays* (Sekhon et al., 2013; Klepikova et al., 2016; Penin et al., 2019).

Using Shannon entropy value lower than 0.25 we identified 521 tissue-specific genes. As in case of direct count the majority of genes was associated with anthers or roots (Fig. 2). According to GO enrichment, genes uniquely expressed in the mature anthers were involved in cell wall organization, biogenesis and modification and had pectinesterase and enzyme inhibitor activity (Table S7). Young anthers were characterized by genes encoding products with amine and amino acid binding activity (Table S8). Root-specific genes (expressed in the sample “Root without apex”) were described by terms “response to chemical stimulus”, “response to oxidative stress”, “oxidation reduction”, and “heme binding” (Table S9).

**Figure 2.**
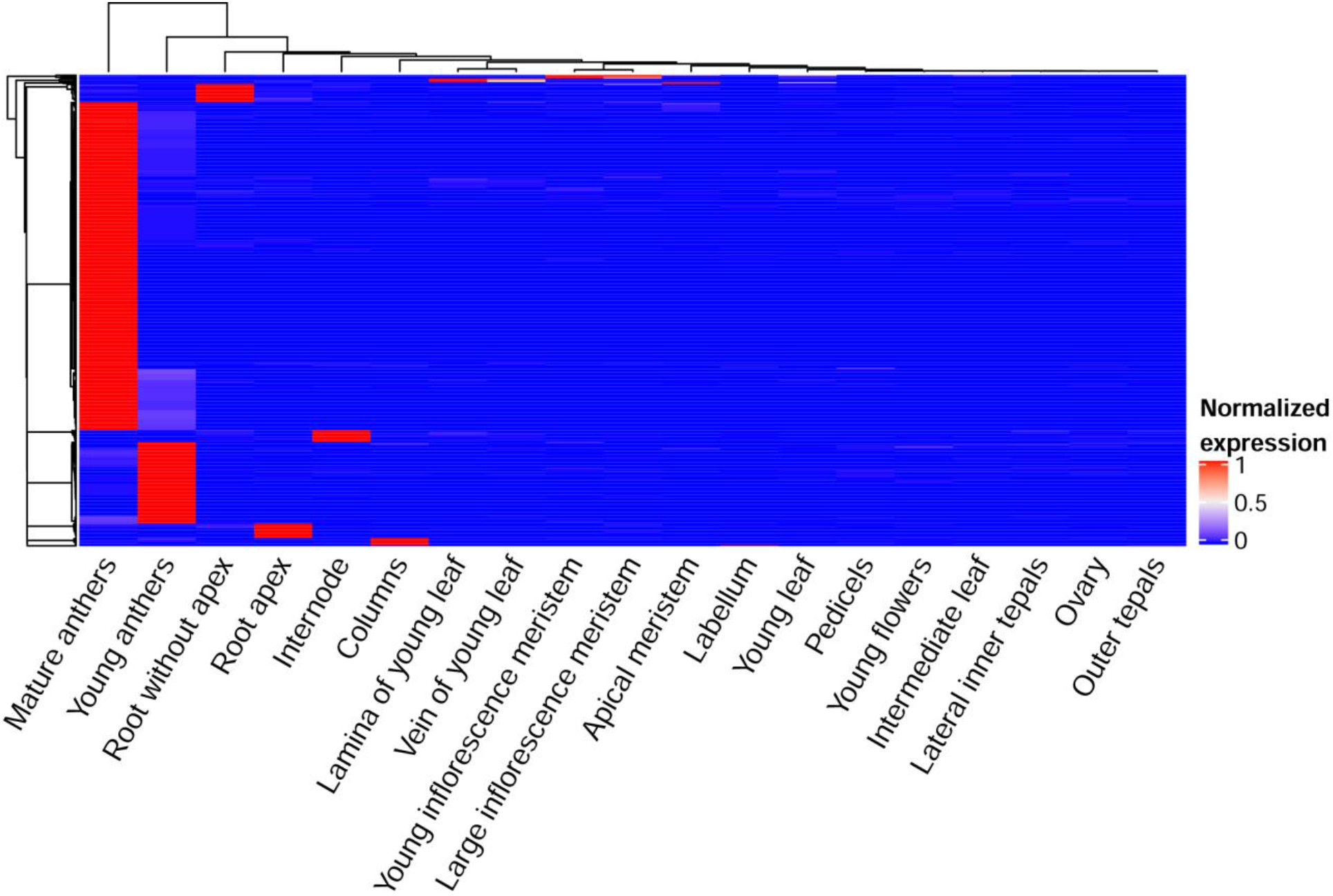
The heatmap of tissue-specific genes. Expression levels of each gene in each sample were normalized on its maximal expression level.

To find genes with opposite behaviour which uniformly expressed across tissues we selected genes with Shannon entropy 3.55 or higher and calculated coefficient of variance (CV) as a measure of expression stability. For 899 out of 1 340 genes CV was less than 0.25, indicating uniform expression in all samples and biological replicates. Stable genes had GO enrichment in terms associated with vesicles, membranes, RNA processing and localization. The list of GO categories strongly overlapped with the enrichment of *A. thaliana* uniformly expressed genes indicating inter-species universality of basic biological processes (Table S10).

### *P. equestris* Transcriptome Variation Database

We aimed to make our transcriptome data easily accessible and ready to use, so we uploaded *P. equestris* transcriptomes into our database Transcriptome Variation Analysis (TraVA, http://travadb.org/browse/Species=Phalaenopsis_equestris/). TraVA interface demonstrates a color chart of gene expression profiles in a single- or multiple-gene view. A user can prefer to show or hide expression values in a chart and choose between several types of read count normalization (Fig. 3).

**Figure 3.**
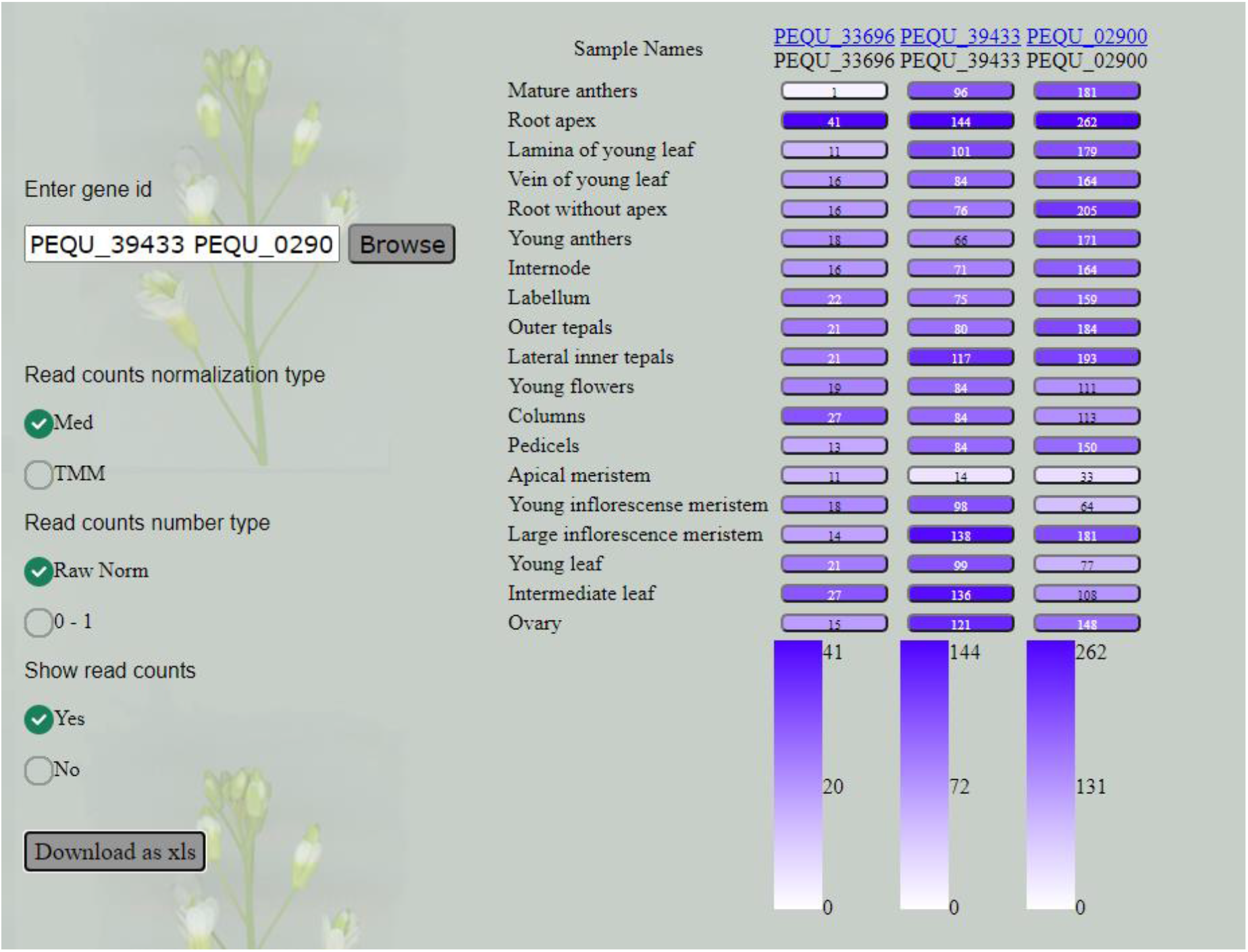
Database view.

### The application of the TraVA database to the characterization of orchid genes

Graphical interface of TraVA facilitate gene expression patterns analysis and comparison and can be widely used in *P. equestris* functional studies. Orchids are a large and highly diverse plant family whose species adapted to a number of ecological niches (typical terrestrial plants, epiphytes, non-photosynthetic plants). These adaptations reflect in their genome – for example, *P. equestris* which has sophisticatedly differentiated perianth the number of *AP3* orthologs is higher compared to *Apostasia schenzhenica*, the basal orchid species with undifferentiated perianth. Vice versa, *P. equestris* which is an epiphyte and does not develop typical terrestrial roots, lacks *AGL12* and several genes of the *ANR1* clade, in contrast to *A. schenzhenica.* This stresses the importance of the study of lineage-specific genes and gene families. For the genes that do not have orthologs in model species the analysis of the expression profiles is the first step towards functional characterization.

We identified 181 (160 after filtering of the proteins that had X on more than 50% of length) *P. equestris* proteins that do not have significant similarity to any *Arabidopsis* protein (e-value cut-off = 10). Out of them 118 share similarity with the proteins of *A. schenzhenica* and are thus presumably orchid-specific while 42 have no hits and thus emerged after the divergence of Apostasioideae and Epidendroideae. The survey of the expression profiles showed that 93 of them are not expressed in any of the samples of the map (Fig. S2). Among the ones which are expressed most are expressed at very low levels. Higher expression levels are associated with reproductive structures, in particular, anthers (Fig. 4).

**Figure 4.**
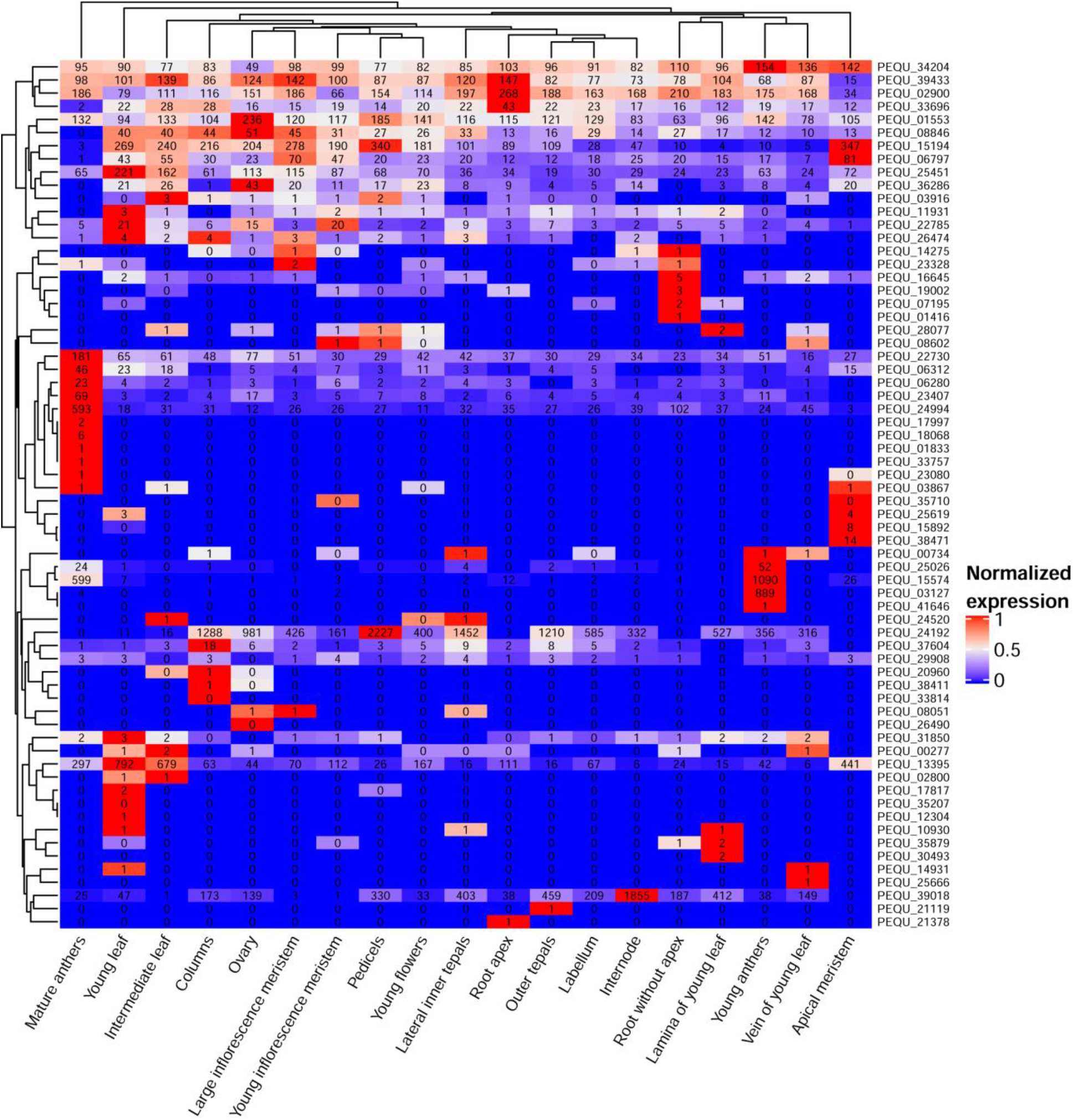
The heatmap of P. equestris-specific genes. Expression levels of each gene in each sample were normalized on its maximal expression level for the color key. The numbers on the figure represent normalized gene read count averaged over biological replicates.

Among vegetative structures the most distinct is root (root apex) where three genes – *PEQU_39433, PEQU_02900, PEQU_33696* – have the highest expression levels. *Phalaenopsis* roots are unique (compared to most other plants, including *A. schenzhenica,* but not to other epiphytic orchids) in many respects – in particular, they are photosynthetic and develop a special structure called velamen. Velamen is a tissue of epidermal origin that consists of several layers of dead cells which help to absorb water and protect photosynthetic tissues of the root from the UV damage. Notably, *PEQU_39433* and *PEQU_33696* do not have homologs in *A. schenzhenica. PEQU_02900* has a marginal similarity (34%) with *A. schenzhenica* protein encoded by Ash001570 gene.

The topic of orphan genes – the ones that lack detectable homologues in other lineages - is widely discussed, in particular in application to plants (Arendsee, Li & Wurtele, 2014). While a part of the orphan genes might represent the artifacts of the annotation, others have a function (for example, *A. thaliana* orphan gene *QQS* which acts in starch metabolism (Li et al., 2009). The functional analysis of orphan genes however lags behind the typical genes; they are overlooked in the annotations based on the homology; they are also usually expressed at lower levels and in a narrower range of tissues (reviewed in Schlötterer, 2015). The study of expression levels and patterns of a potential orphan gene is a first step towards its characterization – the detectable level of expression is an evidence of that the ORF is indeed a gene, not an annotation artifact.

Notably that orphan genes in the well-characterized animal objects (Drosophila, primates) have expression patterns biased towards male reproductive structures (Begun et al., 2007; Xie et al., 2012). According to the “out-of-testis” hypothesis (Kaessmann, 2010) this is mediated by the unique epigenetic state of the chromatin during male gametogenesis. We observed the same bias in *Phalaenopsis*; the growing availability of plant transcriptome maps will enable to find out if this is universal for plants.

## Conclusions

In this study we present a transcriptome map of orchid *Phalaenopsis equestris* covering 19 organs at various stages of the development. We identified 521 tissue-specific genes the majority of which expressed in anthers, roots (11%), and meristems. The uniformly expressed genes were associated with the similar processes as in *Arabidopsis thaliana,* i.e. vesicles, membranes, RNA processing and localization. In order to improve reuse of the data we integrated transcriptome map in our database TraVA and demonstrated its usability in the study of *P. equestris* orphan genes.

## Author Contributions

Conceptualization, A.A.P.; Data curation, A.V.K.; Formal analysis, A.V.K. and A.S.K.; Funding acquisition, A.A.P.; Investigation, A.A.P., M.D.L. and A.V.K.; Methodology, A.A.P and M.A.E.; Project administration, A.A.P.; Software, A.V.K. and A.S.K.; Supervision, A.A.P.; Writing—original draft, A.V.K. and M.D.L..; Writing—review & editing, A.A.P., A.V.K. and M.D.L.

