## Supplementary figures and images for "Transcriptome atlas of *Phalaenopsis equestris*"

### Figure S1

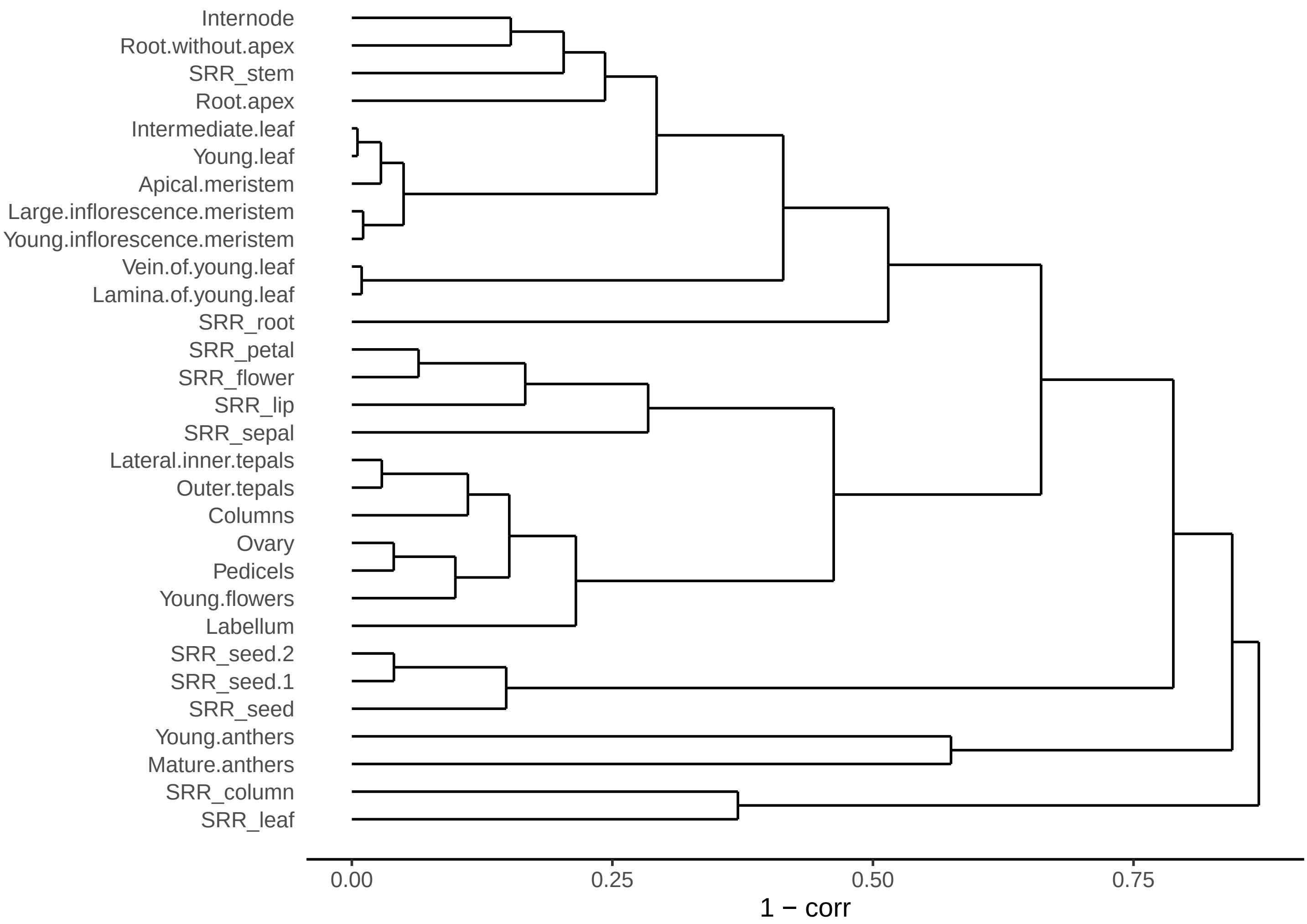

### Figure S2

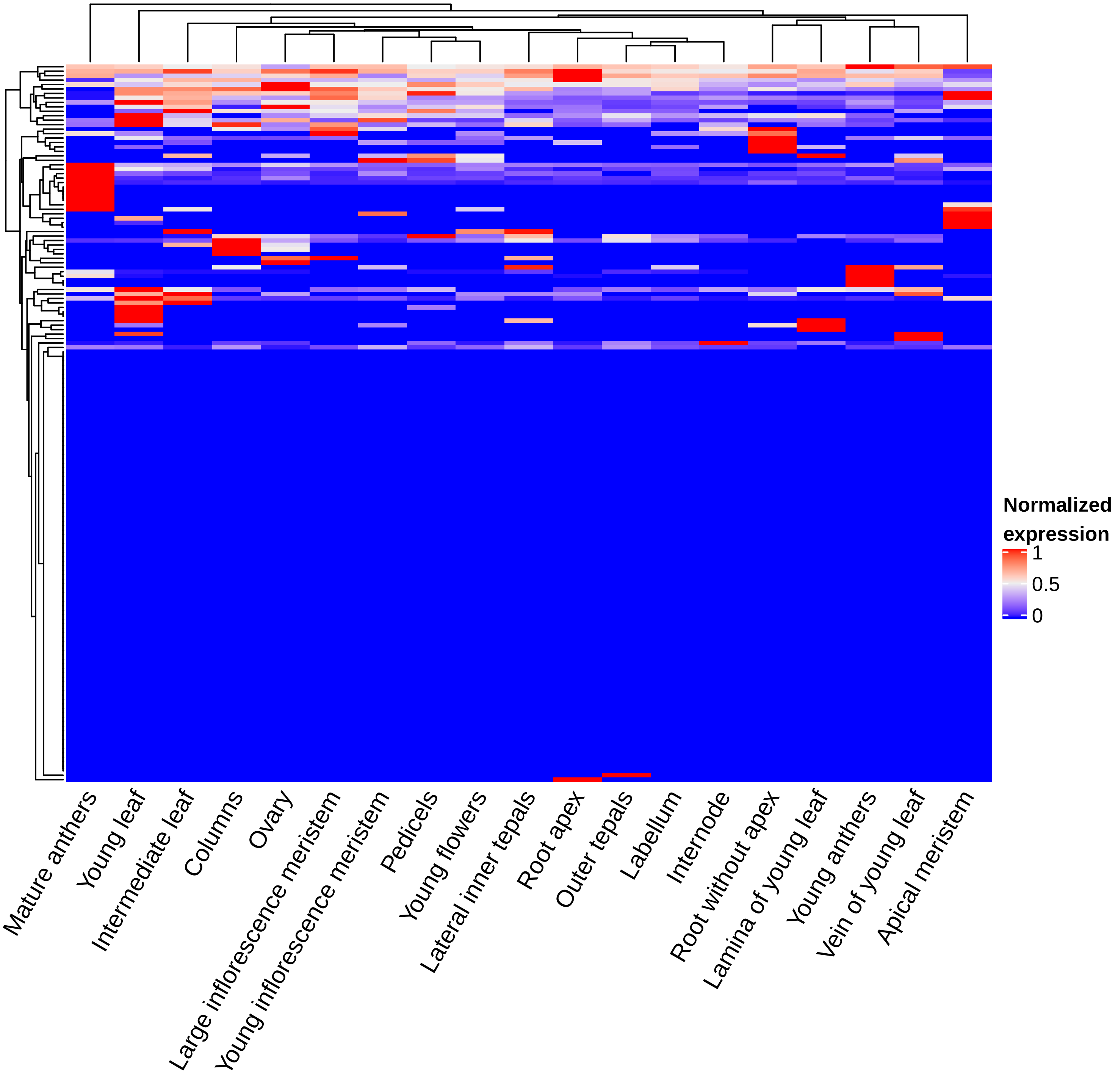
